# Red harvester ants (*Pogonomyrmex barbatus*) do not distinguish between sorghum head mold symptomatic and asymptomatic seeds

**DOI:** 10.1101/2024.07.22.604667

**Authors:** Lilly V. Elliott-Vidaurri, Hannah J. Penn, Robin A. Choudhury

**Affiliations:** Cornell University, Department of Entomology, 2126 Comstock Hall, Ithaca, NY, USA 14853; United States Department of Agriculture, Agricultural Research Service, Sugarcane Research Unit, 5883 Usda Rd., Houma, LA, USA 70360; School of Earth Environment and Marine Sciences, University of Texas Rio Grande Valley, 1201 W University Dr. Edinburg,TX 78539

**Keywords:** Seed preference, fungal infection, ant-seed interactions, Lower Rio Grande Valley

## Abstract

Red harvester ants, *Pogonomyrmex barbatus* (Smith) (Hymenoptera: Formicidae), common in the Lower Rio Grande Valley of Texas, are known to gather seeds from areas around their nests and store the seeds inside their nests for later consumption. As these ants often nest in and near agricultural fields, some of these seeds may be from crops and may also be infected with fungal plant pathogens. These pathogens can degrade seed coats and may cause the seeds to rot within the ant nests, decreasing storage time and potentially spreading the pathogen to other stored seeds. We studied how head mold, a common sorghum disease, changed ant preferences for sorghum seeds. Using seed depots, we evaluated foraging preferences for sorghum seeds with and without head mold and then monitored how many seeds of each type were collected by the colonies after 1, 2, 4, and 24 hours. We found that red harvester ants did not have any significant preference for infected or uninfected seeds, taking both equally over time. Given this non-preference, ants were assumed to be storing infected seeds next to uninfected seeds within their colonies. However, the risk that stored pathogen-infected seeds poses as a source of future seed infection to seeds within the nest and plants in the surrounding field needs to be further examined.

## 1. INTRODUCTION

Part of maintaining healthy colonies in social insects, or social immunity, is preventing and mitigating microbial infection from entering and spreading within the colony (Cremer et al., 2007). Fungal pathogens and associated toxins are often cues that induce social immunity behaviors and physiological responses in social insects such as ants (Liu et al., 2019). One crucial part of preventing pathogens from spreading within a colony is the detection and management of infected food items (Meunier, 2015). This may entail being highly selective of food items to prevent pathogens from entering the nest. For instance, leaf cutter ants preferentially forage on plant materials without certain microbes and then “weed out” unwanted fungal isolates from their fungal garden (Currie & Stuart, 2001; Rocha et al., 2014). Alternatively, management can include processing or cleaning food items to make them more suitable. In the *Atta sexdens rubropilosa* (L.) (Hymenoptera: Formicidae) leaf cutter ants, workers produce mandibular gland secretions that are able to inhibit multiple pathogens, including known plant pathogens, such as *Candida albicans* (de Lima Mendonça et al., 2009), *Botrytis cinerea* (Marsaro Junior et al., 2001), and *Fusarium solani* (Rodrigues et al., 2008). Further, seeds that have been handled by ants and placed in the nest differ in their surface microbial communities (Lash et al., 2020), and some seeds may experience reduced levels of fungal infection after handling by ant workers (Ohkawara & Akino, 2005).

Harvester ants are social insects that gather seeds from the surrounding environments to be stored in the nest for future consumption (MacMahon et al., 2000), and their preferential seed collection may be related to plant species and seed characteristics such as fungal infection status (Elliott-Vidaurri et al., 2022; MacMahon et al., 2000; Penn & Crist, 2018). For instance, a study of *Pogonomyrmex occidentalis* (Cresson) (Hymenoptera: Formicidae) found that foragers collected control seeds from a depot at twice the rate as seeds with external fungal symptoms (Crist & Friese, 1993). Similarly, *P. occidentalis* and *P. rugosus* Emery (Hymenoptera: Formicidae) disproportionately discarded otherwise preferred fescue seeds when those seeds contained endophytic fungi (Knoch et al., 1993). However, seed microbial inoculation status does not always impact selection rates, as previous research on red harvester ants, *P. barbatus* Smith (Hymenoptera: Formicidae), did not find any significant difference in foraging rates for wheatgrass and radish seeds that had been inoculated with a nitrogen-fixing bacteria (Elliott-Vidaurri et al., 2022).

Harvester ants can occur in crop production areas, nesting and foraging near and within the boundaries of commercial crop fields (Uhey & Hofstetter, 2022). Many crops are affected by fungal plant pathogens, particularly pathogens of seed heads that can impact plant health and postharvest storage longevity due to mold (Fleurat-Lessard, 2017). Fungal plant pathogens that affect seeds can sporulate and may spread to other nearby seeds, rapidly contaminating seed stores (Hernandez Nopsa et al., 2015). Some of these fungal pathogens produce secondary metabolites like mycotoxins that can be toxic to insects and other animals even at low doses (Fleurat-Lessard, 2017; Mannaa & Kim, 2017). A major threat to sorghum production in the Lower Rio Grande Valley of Texas is sorghum head mold, caused by multiple causative agents including *Fusarium* spp., *Aspergillus* spp., *Curvularia* spp., *Colletrotrichum* spp., and *Alternaria* spp. (Ackerman et al., 2021). Red harvester ants, *Pogonomyrmex barbatus* (Smith) (Hymenoptera: Formicidae), have been observed locally in grain sorghum fields (Figure 1), a major crop that is susceptible to sorghum head mold. Given the proximity of harvester ants in crop fields with infected seed heads, foraging ants may forage on seeds that have been infected with plant pathogens. Our objective was to determine if harvester ants preferentially forage on symptomatic or asymptomatic crop seeds using seed depots.

**Figure 1.**
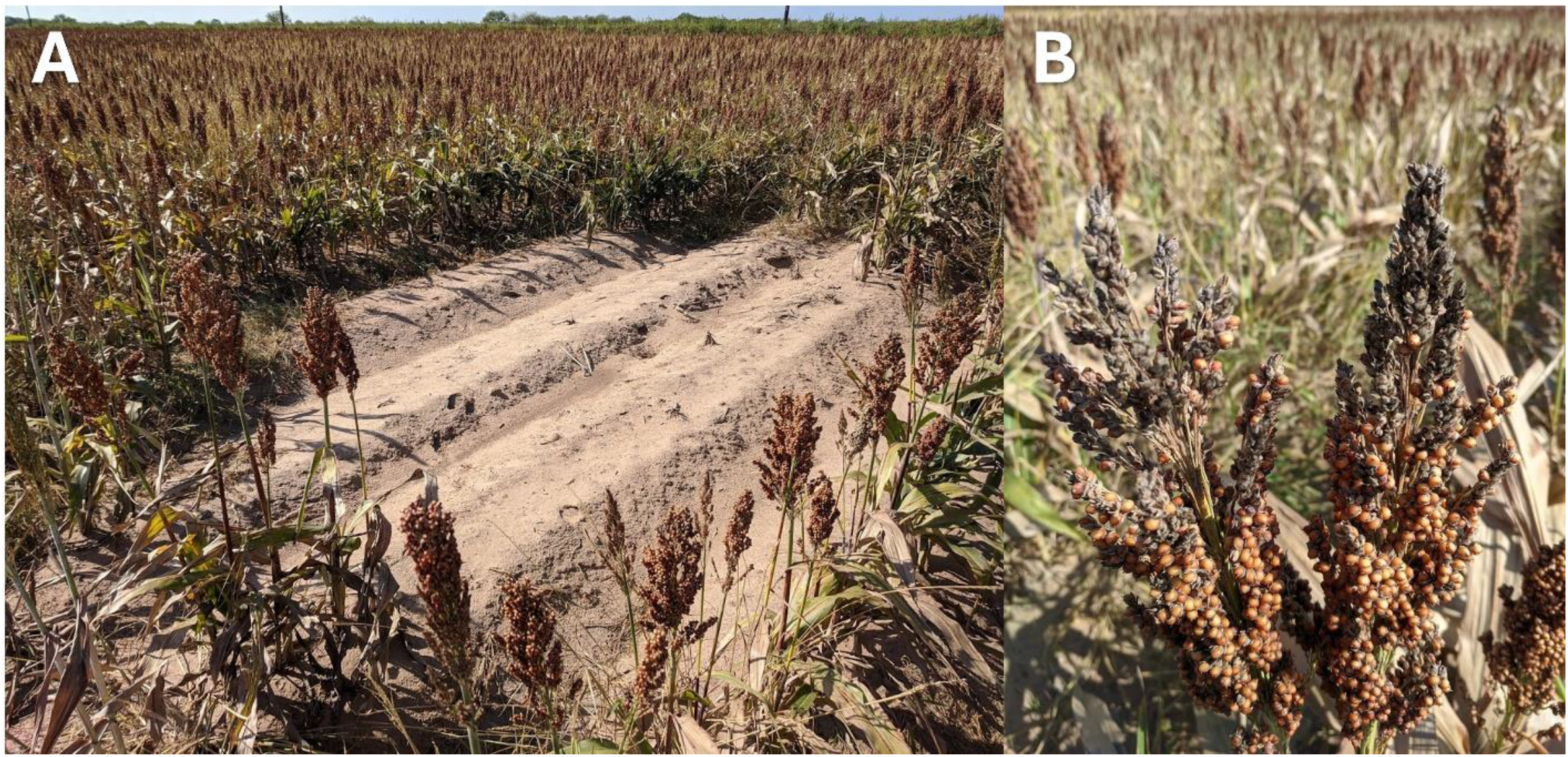
(A) Red harvester ant (*Pogonomyrmex barbatus*) nest disk in a sorghum field in Hidalgo Co., Texas and (B) grain sorghum head displaying symptoms of head mold.

## 2. Materials and Methods

Sorghum seeds (*Sorghum bicolor*) with and without sorghum head mold symptoms were sourced from a commercial grain sorghum field in Monte Alto, TX, USA. Neighboring sorghum fields on the same and two neighboring farms were also briefly observed for red harvester ant presence to ensure natural co-occurrence with infected sorghum seeds. Seeds were then visually sorted into symptomatic and asymptomatic classes then stored at -20 °C until use to prevent further seed degradation.

To determine harvester ant preferences for sorghum seeds with and without symptomatic sorghum head mold, seed depots were deployed according to methods developed in (Elliott-Vidaurri et al., 2022) at 20 active colonies located within the boundaries (∼1.5 km2) of the UTRGV Edinburg, TX campus (26°18’33.1”N 98°10’26.8”W) (Elliott-Vidaurri et al., 2023). This peri-urban location allowed us to evaluate colonies that were naïve to sorghum head mold. All preference tests were completed between March and June 2021 when wind speeds were ≤ 32 km/h on 8-10 colonies at a time that were separated by a minimum of 10 m. Each depot consisted of a sanded I-plate Petri dish, with 3 U-shaped entrance holes in the side for easy seed removal (Elliott-Vidaurri et al., 2022). One side of the plate was marked with a red square to differentiate the treatments. Ten seeds of a single treatment (symptomatic and asymptomatic sorghum) were placed on a randomized side of the Petri dish, with ten seeds of the other treatment on the alternate side. Depots were placed ∼1.8 m along the colony’s main truck trail. Bird and mammal predation was prevented by securing 23×23 cm wire cages made of 1×1 cm hardware cloth around each dish using 4 U-shaped nails. The number of seeds removed was determined after 1, 2, 4 and 24 h; at each timepoint, the temperature, wind speed, and cloud cover percentage were measured.

All statistics were completed in RStudio Version 1.4.1717 (R Core Team, 2023). The Kaplan-Meier survival estimator (function “survfit” in the “survival” package was used to calculate seed removal event likelihood over time and differences between treatments (Therneau et al., 2000). Seeds that were right-censored due to external events (e.g., flipped dishes due to high wind speeds, removal of the cage prior to the 24 hours period, etc.) and those not removed after 24 h were censored.

## 3. Results

Sorghum head mold was observed primarily in several non-irrigated fields throughout Hidalgo and Cameron Counties in Texas, and frequently co-occurred in fields with red harvester ant nests (Fig. 1). When cultured on potato dextrose agar, all tested sorghum seeds that were symptomatic with sorghum head mold were contaminated with fungal species from the genera *Fusarium, Aspergillus, Cladosporium, Epicoccum*, and *Alternaria*, all fungi that are commonly associated with sorghum head mold.

The seed depot trials were run between March and April 2021, with an average temperature of 27.5 °C, an average wind speed of 22.5 kph, and an average cloud cover of 49.3%. Overall, there was no significant effect of seed-health status when assessing the time to removal (p = 0.306), with harvester ants removing both symptomatic versus asymptomatic seeds at relatively similar rates (Fig. 2). Both treatments had roughly half of their seeds removed within the first 4 h and all seeds from both treatments were removed after 24 h in all tested sites.

**Figure 2.**
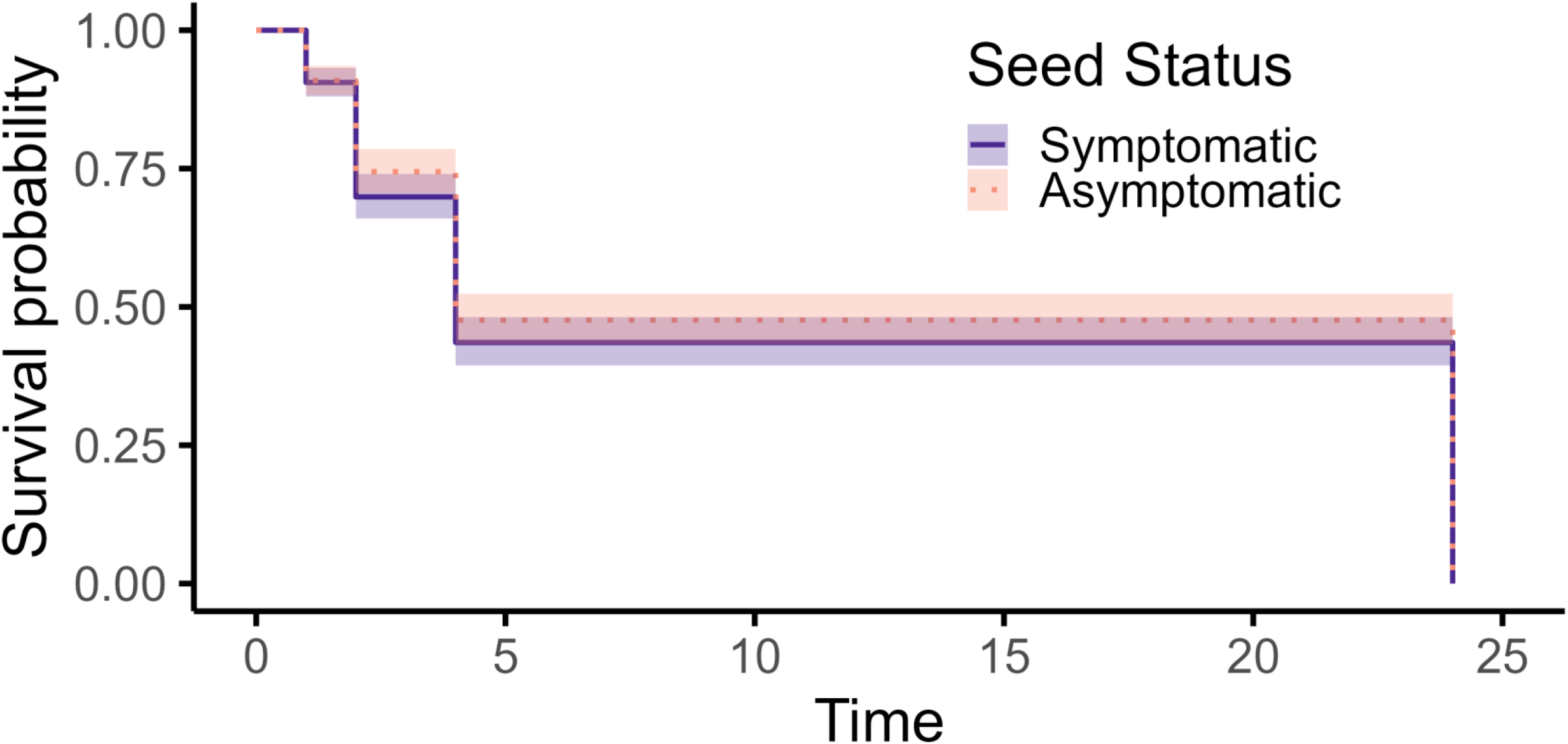
Kaplan-Meier curves (n = 20) indicating the likelihood of seed removal [survival] by red harvester ants (Pogonomyrmex barbatus) over the course of the seed preference trial (observations at 1, 2, 4, and 24 h) for sorghum seeds symptomatic or asymptomatic for sorghum head mold.

## 4. Discussion

Although prior work has indicated that several species of harvester ants often reduce foraging or increase removal of seeds containing fungal infection (Crist & Friese, 1993; Knoch et al., 1993), we did not observe any differences in the foraging by P. barbatus on sorghum seeds with and without head mold symptoms. However, we were unable to determine the fate of collected seeds. In some cases, harvester ants may initially collect seeds and then drop them before storage in the nest or take the seeds into the nest before later removing them to an external refuse pile (Arnan et al., 2010; MacMahon et al., 2000; Mull, 2003). Further, we only observed these interactions during a portion of the year, and changing environmental conditions may alter foraging selectivity by limiting food availability and/or increasing environmental stress (Whitford & Ettershank, 1975). Our use of naïve ant colonies helped to prevent bias in our experiments, however ants that had previously interacted with symptomatic seeds may experience seed learning and memory (Johnson, 1991), altering the likelihood of future interactions. Future studies should explore how past experiences with symptomatic seeds could influence the likelihood of seed selection.

Regardless of final seed fate, ant dispersal of infected seed even partially across a crop field may facilitate plant disease spread. However, if the infected seeds were stored for any length of time within the nest, then potentially large numbers of diseased seeds may be held underground, allowing pathogens to persist between cropping years. This becomes a greater concern to growers if the nest granary is shallow enough to be disrupted when the field is tilled during standard agricultural management. Prior observations of this species indicate that granaries can be fairly close to the soil surface (4-75 cm deep), with whole seeds in the upper galleries and shucked seeds in lower galleries and some galleries being filled with seed covered in a “glutinous material” (McCook, 1880). Tillage can reach a depth of 25 cm, meaning that the granaries of P. barbatus may be potential sources of both seed and disease inoculum in agricultural fields. However, the viability of plant pathogens through this dispersal pathway needs to be further evaluated.

Although plant disease symptoms did not alter the removal rate in our study, many ant species can detect and alter the microbial communities in their immediate surrounding environments (Christe et al., 2003; Lindström et al., 2019). For instance, active *P. occidentalis* nests were associated with an increased density of mycorrhizal fungi in the surrounding soil (Friese & Allen, 1993). This may be directly through their manipulation of seeds (Ohkawara & Akino, 2005) and other plant materials (Christe et al., 2003) and secretion of antimicrobial materials (de Lima Mendonça et al., 2009; Marsaro Junior et al., 2001; Rodrigues et al., 2008). Or these changes may be mediated more indirectly by changing soil properties and chemistry (Wagner et al., 1997) or plant communities (Whitford & Ettershank, 1975). While some ants avoid soil with higher densities of entomopathogenic fungi (Huang et al., 2020), other ant genera have also been shown to not avoid foraging in areas even if entomopathogenic fungi are present in the environment (Pereira et al., 2021). Ultimately, understanding how ants interact with a broader range of microbial partners can expand our understanding of their role in agroecological processes in understudied regions.

## Acknowledgements

We would like to acknowledge Joseph Rabago, Daniella Rivera, and Adrian Noval for technical assistance in this project. We would like to thank Dr. A. Wright for manuscript comments. Mention of trade names or commercial products in this publication is solely for the purpose of providing specific information and does not imply recommendation or endorsement by the U.S. Department of Agriculture. USDA is an equal opportunity provider and employer.

## Author contributions

All authors contributed to the study conception and design. Material preparation and data collection were performed by L.V.E.-V. Statistical analyses and figure development were performed by L.V.E.-V. The first draft of the manuscript was written by H.J.P. and R.A.C. and all authors commented on previous versions of the manuscript. All authors read and approved the final manuscript.

